# Effect of disease prevalence and growth stage on symptoms severity in the *Turnip mosaic virus - Arabidopsis thaliana* pathosystem

**DOI:** 10.1101/2023.04.12.536568

**Authors:** Francisca de la Iglesia, Santiago F. Elena

## Abstract

Plants emit volatile organic compounds (VOCs) in response to biotic and abiotic stimuli that provide information about their physiological status to other individuals in the community. Nearby receivers adjust their own defenses in response to these chemical cues. The majority of studies to date has concentrated on the communication of abiotic stressors (*e.g*. salinity or drought) or herbivory. Less attention had received the role of VOCs during microbial infections and almost nothing has been done for viruses. Here we investigated the function of VOCs during turnip mosaic virus infection of *Arabidopsis thaliana*. First, we looked at the influence of two factors on the kinetics of symptoms progression in receivers, namely the prevalence of infection in the population and the growth stage of the receiver plants at inoculation. We found that young plants were more sensitive to the protective effect of VOCs than older ones, and that high infection prevalence results in a slower disease progression in receivers. Second, we tested the possibility that jasmonates could be VOC candidates. To do this, we examined the kinetics of symptoms progression in jasmonate-insensitive and wild-type plants, and the results showed that the protective effect vanished in the mutant plants. Third, we investigated the possibility that root communication would be also relevant. We found that the kinetics of symptom progression across receivers was further slowed down in an age-dependent manner when plants were planted in the same pot. Together, these preliminary findings point to a potential function for disease prevalence in plant communities in regulating the severity of symptoms, this effect being mediated by VOCs.

---

To protect themselves from biotic and abiotic stresses, plants create and emit a wide variety of volatile organic compounds (VOCs) (Bitas et al. 2013; Brosset and Blande 2022; Loreto and D’Auria 2022). These VOCs may also serve as warning signals to prime defenses in unaffected areas of the same plant and in neighboring plants. Plant-to-plant communication is a phenomenon that was first identified by Baldwin and Schultz (1983). Since this groundbreaking research, a plethora of investigations have provided strong experimental support for the hypothesis that plants release VOCs in response to a variety of stresses, like salinity (Lee and Seo 2014), drought (Falik et al. 2022), herbivory (Dolch and Tscharntke 2000), infection by bacterial (Riedlmeier *et al*. 2017), fungal (Moreira *et al*. 2020) or viral (Ghosh *et al*. 2022) pathogens, or even triggering plant-plant allelopathy to minimize intraspecific competition (Santonja *et al*. 2019).

The chemical nature of VOCs is diverse and, indeed, VOCs typically act as complex blends of various volatiles rather than as single molecular species (Ueda *et al*. 2012). Among the most frequently described ones are terpenoids, oxylipins [most notably jasmonic acid (JA) and some of its derivatives, such as methyl-JA and *cis*-JA], olefins (*e*.*g*, isoprene and ethylene), and methylsalicylate (MeSA). The phenylpropanoid pathway, which produces the volatile MeSA, and the methyl erythritol phosphate pathway, which produces volatile isoprenoids like monoterpens and hemiterpens, are examples of biochemical pathways induced by abiotic and biotic stresses and producing VOC emissions that may be used as signals or to prime defenses (Loreto and D’Auria 2022).

VOCs modulate JA signaling involved in the induction of defenses, and it is mostly assumed that by enhancing defenses, VOCs would inhibit the growth of receivers if investment in defense trades off with growth and development. Indeed, several studies support the notion that exposure to VOCs affect growth and reproduction to an extent that varies among plant species (reviewed in Brosset and Blande 2022).

The impact of plant-to-plant communication on virus epidemiology is unknown. Given that infected plants emit VOCs, it stands to reason to assume that as epidemics spread through a population of vulnerable hosts and the incidence of infection rises, the concentration of VOCs will rise as well. This will prime antiviral defenses in receiver non-infected plants more effectively, increasing tolerance to infection and slowing the progression of symptoms in primed plants. Furthermore, it is anticipated that plants at different developmental stages would respond to this priming in different ways, with younger plants being more sensitive to VOCs than older ones that are already investing in reproduction. To test these two hypotheses, we have used the turnip mosaic virus (TuMV) - *Arabidopsis thaliana* natural pathosystem. In a first set of experiments, we will test whether the kinetics of disease progression (*i*.*e*, the severity of symptoms) of receiver plants was affected by the prevalence of infection in the population (*i*.*e*, the frequency of infected plants). Then, we will test whether the effect was dependent on the developmental stage of the receivers and duration of exposition. In a second set of experiments, we will test the potential role of JA as chemical primer of increased tolerance to infection. Finally, we will show results of a third experiment testing whether root communication may also contribute to priming a more tolerant state in receiver plants.

## Materials and Methods

### Plants and growth conditions

*Arabidopsis thaliana* (L.) HEYNH of the accession Col-0 were maintained in a BSL2 climatic chamber under a photoperiod of 8 h light (LED tubes at PAR 90 - 100 μmol m−2 s−1) at 24 °C and 16 h dark at 20 °C and 40% relative humidity.

Mutant *jasmonate insensitive 1* (*jin1*) (AT1G32640) was used to test the potential role of jasmonates as VOCs in signaling viral infection. Mutant *jin1* exhibits decreased induction of JA-responsive genes, including *PR1*. However, mutant plants show reduced sensitivity to *Pseudomonas syringae* and *Botrytis cinerea* infections (Nickstadt *et al*. 2004; Laurie-Berry *et al*. 2006). Indeed, Nickstadt *et al*. (2004) showed that *jin1* was defective in JA sensing but not in jasmonates biosynthesis, and that infections were associated with two-fold increases in salicylic acid (SA), suggesting a possible contribution of the SA pathway to *jin1* plants resistance to the above-mentioned pathogens. This would suggest that over-expression of one pathway does not necessarily down-regulates the other (Nickstadt *et al*. 2004). Furthermore, these authors concluded that *JIN1* affected basal resistance rather than resistance based on gene-for-gene interactions. In general, enhanced JA accumulation favors plant defense against viral infections (Islam *et al*. 2019); and for example, *jin1* plants show increased susceptibility to TuMV infection (Navarro *et al*. 2022).

### Virus, inoculation procedure and characterization of symptoms

TuMV (species *Turnip mosaic virus*, genus *Potyvirus*, family *Potyviridae*) infectious sap was obtained from TuMV-infected *Nicotiana benthamiana* DOMIN plants inoculated with the infectious plasmid p35STunos containing a cDNA of the TuMV genome (GeneBank accession AF530055.2) under the control of the cauliflower mosaic virus 35S promoter and the *nos* terminator (Chen *et al*. 2003) as described elsewhere (González *et al*. 2019; Corrêa *et al*. 2020). This TuMV sequence variant corresponds to the YC5 strain from calla lily (*Zantesdeschia* sp.) (Chen *et al*. 2003). After plants showed symptoms of infection, they were pooled and frozen with liquid N_2_. This frozen plant tissue was homogenized into a fine powder using a Mixer Mill MM400 (Retsch GmbH, Haan, Germany).

For inoculation of *A. thaliana* plants, 0.1 g of powder were diluted in 1 mL of inoculation buffer (50 mM phosphate buffer pH 7.0, 3% PEG6000, 10% Carborundum) and 5 μL of the inoculum were gently rubbed onto three leaves per plant. Depending on the particular experiments, plants were inoculated at either of the following two growth stages (Boyes *et al*. 2001): (*i*) at stage 3.32, when the rosette is ∼25% of its final size; (*ii*) at stage 5.10, the precise moment at which the first flower buds are visible, marking the transition from juvenile vegetative growth to adult flower development. Hereafter we will refer to plants at growth stage 3.32 as *prebolting* and to plants at growth stage 5.10 as *bolting*.

Severity of symptoms was evaluated in the semiquantitative discrete scale shown in Fig. 1 of Butković *et al*. (2021). This scale ranges from zero for non-infected and asymptomatic plants to five for plants showing a generalized necrosis and wilting. Symptoms severity were annotated daily for each individual plant from inoculation up to 14 days post-inoculation (dpi).

**Fig. 1.**
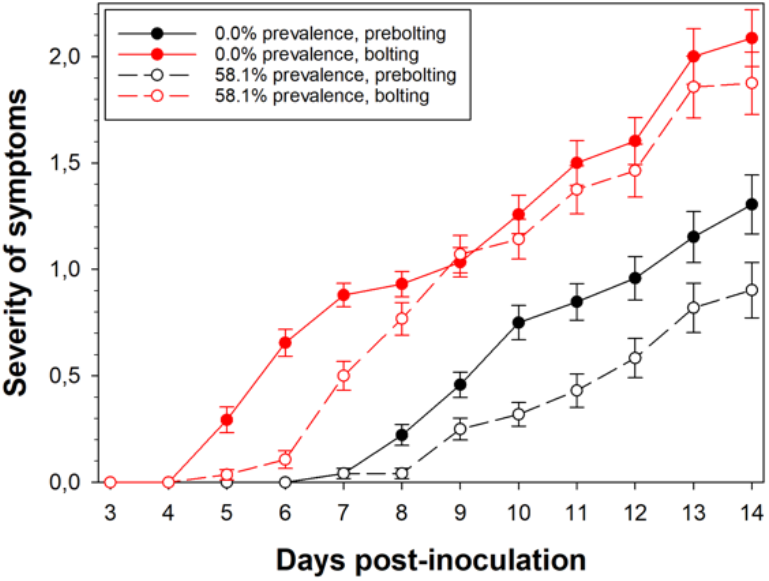
Symptoms progression curves at low and high prevalence of infection and considering the plant growth stage (3.32 as prebolting and 5.10 as bolting) at the time of inoculation. Error bars represent ±1 SEM.

### Evaluating the role of disease prevalence and growth stage in symptoms progression

The interaction between the growth stage of the inoculated plants and the prevalence of disease in the plant community was investigated in the first experiment. There were two distinct prevalence scenarios simulated: the first was zero prevalence, in which no plants in the community were infected during the growing of the 130 wild-type (WT) receiver plants to be infected once reached the desired growth stage. Second, high prevalence, in which the median prevalence of infection among emitters during the development of an additional set of 128 WT receiver plants was 0.318 ±0.109 (±IQR). After inoculation of the receivers at the predetermined growth stage, the median prevalence increased to 0.581 ±0.303. For both prevalence values, 72 plants were inoculated when they reached prebolting while the remaining 56 were inoculated at bolting. The zero and high prevalence treatments were done in two separated growth chambers.

In the second experiment, meant to examine the role of JA as VOC in this pathosystem, WT control and *jin1* plants were all inoculated at prebolting in two prevalence conditions: (*i*) zero prevalence, with 86 WT and 86 *jin1* plants inoculated at the same time. (*ii*) A prevalence of 0.662 ±0.034 among WT emitters during the vegetative growth of the receivers, rising up to 0.974 ±0.107 after inoculation of 80 WT and 72 *jin1* receivers.

### Evaluating the role of root-mediated communication

A third experiment was designed to test the possible contribution of root communication. Two treatments were done, all using WT plants. First, sowing two plants per pot to allow for possible additional root communication. Second, sowing only one plant per pot thus blocking root communication. In both treatments, the distance among plants was kept constant. For the same pot treatment, 130 emitters, one per pot, were inoculated at prebolting. Eighty-two receivers were inoculated the same day and 48 more 12 days later (thus at bolting) sharing a pot with the emitters.

For the different pot treatment, the inoculation design was as follows: 64 emitters were inoculated at prebolting and, the same day, 40 receivers were also inoculated. Twelve days later 24 plants were inoculated at bolting.

### Statistical analyses

To examine the kinetics of disease severity progression, two different tests were conducted. To assess overall differences between curves, nonparametric Wilcoxon signed-rank tests (Wilcoxon 1945) were performed. Second, daily severity data were fitted to repeated measures ANOVA models (Girden 1992) in which days-post inoculation was treated as the within-plant factor (repeated measures) whereas other between-plant factors were treated as orthogonal. In all these models, the type III sum of squares were used to partition total variance among factors and the Greenhouse-Geisser assumed sphericity was used for the within-plant tests (Greenhouse and Geisser 1959). The power of each test, 1 – *β* has also been computed, whereβ is the type II error. Cohen’s 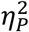 statistic, which gauges the percentage of total variability attributable to each model factor, was used to assess the effect size of the different factors in the ANOVA models (Cohen 1973). Values of 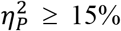 are typically regarded as large effects.

All these statistical analyses were done with SPSS version 28.0.1.1 (IBM, Armonk NY, USA).

### Data availability

Three Excel files containing the disease severity raw data from each experiment described above are available at zenodo.org under doi: 10.5281/zenodo.7663956.

## Results

### The kinetics of symptom development are determined by disease prevalence among emitters and the receivers’ growth stages in an additive manner

Our first set of experiments evaluated the combined contribution of two factors on the progression of disease symptoms: the prevalence of disease among emitter plants and the growth stage of the receiver plants. Fig. 1 shows the disease severity progression curves for two prevalence values and two growth stages for inoculated receiver plants. In both cases dynamics of symptoms development are slower if receivers grew in a plant community with high incidence of infection. When receiver plants were inoculated at prebolting, symptoms started appearing 7 dpi, being on median 19.44% ±38.20 (±IQR) stronger for receiver plants grown in a community with zero prevalence of infection during their development (Fig. 1, black symbols; Wilcoxon signed-rank test: *z* = –2.3664, *P* = 0.0180). However, when receiver plants were infected at bolting, symptoms started being visible 5 dpi, also being on median 14.10% ±13.60 more severe in the case of receivers grown in a community with zero prevalence (Fig. 1, red symbols; *z* = –2.7011, *P* = 0.0069).

To further understand the source of statistical differences, we fitted the data in Fig. 1 to a repeated-measures ANOVA (Table S1) with the degree of prevalence and growth stage as random between-plant orthogonal factors. Obviously, the most significant and of larger magnitude effect corresponds to differences among days (*P* < 0.0001. 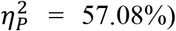). But three other interesting conclusions can be drawn from Table S1: (*i*) Prevalence has significant effects both within- and between-plants (*P* = 0.0442 and *P* = 0.0006, respectively) although of small magnitude 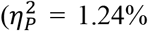 and 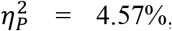 respectively). This confirms that receiver plants grown at high prevalence of TuMV infection develop weaker symptoms (compare solid *vs* open symbols in Fig. 1). (*ii*) More severe symptoms were developed by older than by younger plants, being this effect highly significant both within- and between-plants (*P* < 0.0001 in both cases), although the effect was of moderate magnitude in the within-plant test 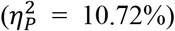 but very large in the among-plants test 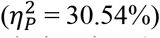 (compare red *vs* black symbols in Fig. 1). (*iii*) Finally, the test of the interaction between prevalence and growth stage was significant in the within-plants test (*P* = 0.0050), although of small effect 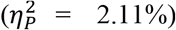 but not significant in the between-plants test (*P* = 0.8104). These tests suggest a similar overall effect of prevalence at both growth stages (compare the difference between solid and open red symbols *vs* the difference solid and open black symbols in Fig. 1).

### Jasmonate as candidates for signaling VOCs

After observing a significant effect of disease prevalence among emitters in the kinetics of symptoms development of receivers, we sought to explore the potential role of JA as a candidate VOC regulating the response of receptor plants. The choice of this hormone was done based in its well stablished role in plant defense against viral infections (Islam *et al*. 2019). Keeping now constant the growth stage at prebolting and varying only disease prevalence, we inoculated WT and *jin1* receiver plants. The results of this experiment are shown in Fig. 2. As in the previous experiment, a highly significant effect was observed in WT plants; in median, plants grown at high TuMV prevalence showed 27.76% ±32.89 weaker symptoms (black symbols in Fig. 2: *z* =– 2.9341, *P* = 0.0033). However, the effect vanished in *jin1* plants, being symptoms progression curves overlapping for both prevalences (red symbols in Fig. 2: *z* =–0.9780, *P* = 0.3281), supporting the role of JA as VOC.

**Fig. 2.**
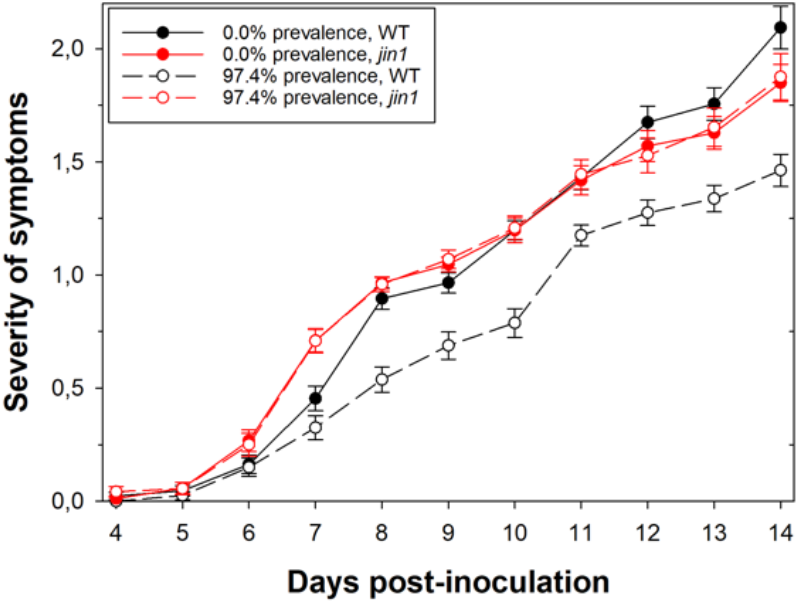
Jasmonate insensitive (*jin1*) plants do not show reduced susceptibility to infection when grown at high prevalence of infection. Error bars represent ±1 SEM.

To gain further statistical insights, we fitted the data in Fig. 2 to a repeated-measures ANOVA (Table S2) with degree of disease prevalence values and plant genotype as random between-plant orthogonal factors. Again, the largest amount of observed variability was explained by differences among days (*P* < 0.0001, 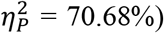. Regarding the three other factors in the model: (*i*) confirming the previous results, prevalence *per se* has a highly significant effect both within- and between-plants (*P* < 0.0001 in both cases), although it was of very large magnitude within-plants 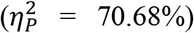 but small between-plants 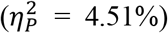 (*ii*) Highly significant differences exist between the symptoms developed by WT and *jin1* plants both when testing within-(*P* = 0.0020) and between-plants (*P* < 0.0001) effects, although both were of small magnitude 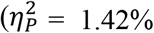 and 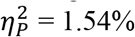, respectively).

And, (*iii*) a highly significant genotype-dependent effect of disease prevalence has been observed both within-(*P* = 0.0013) and between-plants (*P* < 0.0001). In both tests, the magnitude of the effects was rather small 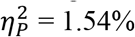 and 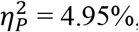 respectively).

### Root contact between emitters and receivers slows the rate at which bolting, but not prebolting, receivers develop symptoms

In our last investigation, we explored the possibility that the TuMV - *A. thaliana* pathosystem may possibly be influenced by root chemical communication between infected and non-infected plants. To verify this idea, seeds were sown singly in pots or in pairs, always keeping the same spacing between them. First, at prebolting, emitters and receivers were inoculated concurrently. In a subsequent experiment, half the plants (emitters) were inoculated at prebolting while the other half were at growth stage 5.12. (receivers). Fig. 3 summarizes the results from these experiments. When emitters and receivers were in separated pots, the growth stage effect described above holds (*z* =– 2.2258, *P* = 0.0262), with receiver older plants developing 15.12% ±15.61 median stronger symptoms than younger ones. However, if plants were potted in pairs, symptoms were on median 28.27% ±11.31 weaker in the receiver older plants (*z* =–3.3510, *P* = 0.0008), a result that sharply contrast with all previous observations and that suggests a possible role for root communication.

**Fig. 3.**
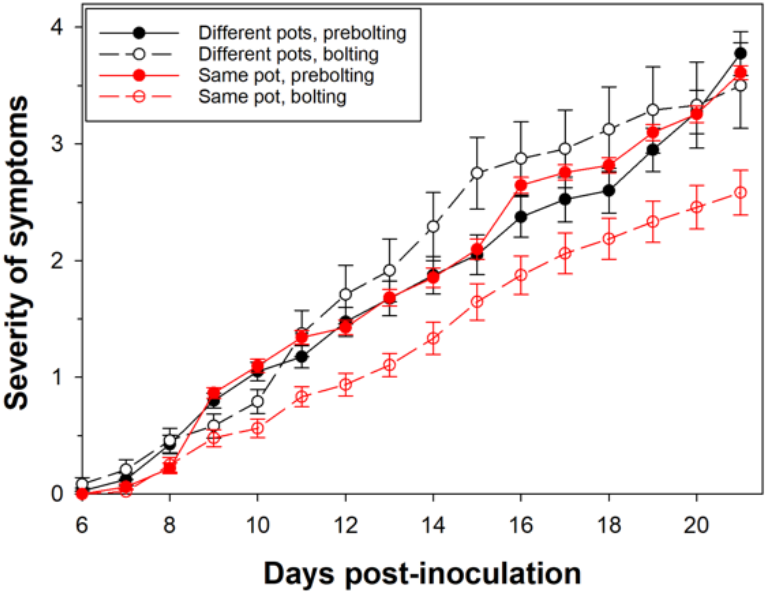
Effect of emitter and receiver plants sharing or not the same pot in symptoms progression curves. Differences among prebolting (3.32) and bolting (5.10) growth stages have been tested. Error bars represent ±1 SEM.

Once more, we fitted the data in Fig. 3 to a repeated-measures ANOVA with two between-plants orthogonal factors: difference in growth stage among emitters and receivers and whether the plants were in the same or different pots (Table S3). Focusing first in the within-plants effects, the largest amount of variability was explained by the progression of symptoms severity along time (*P* < 0.0001, 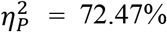). Significant overall effects within-(*P* = 0.0129, 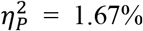) and between-plants (*P* = 0.0022, 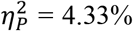) have been found for the number of plants per pot, although the small magnitude. Also, significant overall effects have been associated to the age of the receivers at the time of their inoculation, both within-(*P* = 0.0002, 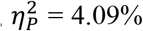) and between-plants (*P* = 0.0053, 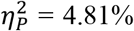), although again, the magnitude of these effects is also small. Finally, the interaction between these two factors is also highly significant, both within- (*P* = 0.0006, 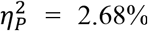 and between-plants (*P* = 0.0005, 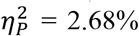). This significant interaction, although certainly of small magnitude, supports that the growth-stage dependent effect on symptoms severity strongly depends on whether plants were able of root communication or not, being weaker in older plants sharing a pot with an emitter during development.

## Discussion

Plants release VOCs in response to biotic and abiotic stimuli, as has been well established. These VOCs represent chemical information on the physiological status of the emitter plants that is sensed by the receiver plants. In response, receiver plants activate defense mechanisms to ensure their homeostasis (Peñuelas and Llusià 2004; Ueda *et al*. 2012; Brosset and Blande 2021; Loreto and D’Auria 2022). In particular for cultivated systems, plant diseases are a significant source of stress. Several studies have documented how infectious diseases affect plant VOCs (Hammerbacher *et al*. 2019; Moreira *et al*. 2020). Although the majority of research on VOCs and infection has been done on bacterial and fungal pathogens, viruses have also been studied (reviewed in Hammerbacher *et al*. 2019). For example, Shulaev *et al*. (1997) showed that increased emissions of MeSA by *Nicotiana tabacum* plants infected with tobacco mosaic virus (TMV) boosted resistance against this pathogen in close-by healthy plants.

In this really preliminary study, we focused on four main questions that might exert an impact in virus epidemiology and evolution: (*i*) in *A. thaliana* communities with varying levels of TuMV prevalence, do VOCs play a role in signaling infection among plants? (*ii*) Does the receiver plants’ age affect how they react to VOCs? (*iii*) Do jasmonates function as VOCs to alert *A. thaliana* populations to TuMV infection? And (*iv*) do root exudates aid in the dissemination of disease information at the community level?

### Social communication and disease prevalence

Our findings suggest a negative correlation between the speed of disease progression and the prevalence of TuMV infection in the plant community. VOCs are effective messengers for stablishing a communication network among plants. However, the potential epidemiological implications of this communication networks have not been considered yet. In recent years, theoretical epidemiologists have been examining the relationship between the propagation of epidemic awareness and the actual epidemic infection in spatially-structured populations of susceptible hosts (Granell *et al*. 2013; Wang *et al*. 2015; Scatà *et al*. 2016; Li and Xiao 2021; Gao *et al*. 2022). In such models, the spread of the epidemics is not anymore properly described by classic models (*e*.*g*, susceptible - infected - recovered, SIR) but more complex and richer dynamics emerge, in which the onset of the epidemics has a critical value defined by the awareness dynamics. In the particular case of plant disease epidemics, such awareness dynamics would depend *e*.*g*. on the gradients of spatial spread of VOCs across the community. Theoretically, the efficient activation of defense mechanisms in receivers, with a concomitant reduction in the rate of symptoms development and virus accumulation are expected to contain the spread of the disease at long distances. Clearly, this possibility deserves further theoretical and experimental work.

### Age-dependent susceptibility to VOCs and the severity of symptoms

Related to our second question, in a recent study, our group has shown that the severity of symptoms associated to TuMV infection of *A. thaliana* actually depends on the growth stage of the plant at the time of inoculation, with older plants developing stronger symptoms (Melero *et al*. 2023). This observation is extensible to viruses from other families and host species (Huang *et al*. 2020; Melero *et al*. 2023). Here, we have verified this earlier finding and shown that the magnitude of the observed effect was unrelated to the prevalence of the disease in the population: older plants systematically show stronger symptoms and are less sensitive to VOCs than younger ones.

Although the defense mechanism known as age-related resistance (ARR) appears to be widespread in plants, it has primarily been studied in the context of plant-bacterial interactions (Kus et al. 2002; Hu and Yang 2019). Our findings, which concur with those made by others (Huang *et al*. 2020; Melero *et al*. 2023), appear to be at odds with ARR. There are instances where older plants are more resistant to viral infections than younger plants, but other species exhibit the opposite pattern. In the cucumber mosaic virus (CMV) - *Capsicum annuum* pathosystem, García-Ruiz and Murphy (2001) demonstrated that the severity of symptoms and viral accumulation were lower in older plants. Similarly, Levy and Lapidot (2008) demonstrated that plant age had little bearing on the severity of symptoms in the tomato yellow leaf curl virus - *Solanum lycopersicum* pathosystem, despite juvenile plants having much reduced fruit output. In stark contrast, Huang *et al*. (2020) discovered that the tomato spotted wilt virus - *A. thaliana* and CMV - *A. thaliana* pathosystems had a developmentally regulated rise in susceptibility, with older plants being more susceptible and exhibiting more severe symptoms.

In the contemporary context, when anthropogenic activities and climate change are changing plant developmental patterns and timing, the influence of the host growth stage on its vulnerability to viruses is particularly pertinent. Abiotic factors connected to climate change have been shown to have genotype-dependent effects on the induction and retardation of developmental processes, including blooming (Craufurd and Wheeler 2009; Tun *et al*. 2021). Moreover, it has been noted that human activities like urbanization (Neil and Wu, 2006) or biodiversity loss (Wolf *et al*. 2017) can influence flowering timing. Viral populations will be exposed to host populations with altered levels of sensitivity as a result of these changes in plant growth. One crucial component to enhancing the management of viral infections in the future is understanding how these developmental changes may affect the evolution of viruses.

### Jasmonate as a candidate for VOC in the TuMV - *A. thaliana* pathosystem

Regarding our third question, although VOCs are composed by a complex mixture of molecules (Brossete and Blance 2021; Loreto and D’Auria 2022), jasmonates (*e*.*g*, methyl jasmonate) have been recognized before as essential airborne signals generated by the emitters and inducing defense responses in the receivers (Tamogami *et al*. 2008). Furthermore, the role of JA in defense responses against viral infections has been well stablished (Islam *et al*. 2019). To test such role of JA in the TuMV - *A. thaliana* pathosystem we used *jin1* mutants, which are insensitive to JA, as receivers. In good agreement with the predicted role of jasmonates, *jin1* plants grown in a community with high TuMV prevalence did not show any delay in symptoms progression compared to WT plants after inoculation. Other studies had suggested a similar role for MeSA in the TMV - *N. tabacum* pathosystem (Shulaev *et al*. 1997). JA and SA signaling pathways are thought to be engaged in an antagonistic tradeoff (Glazebrook 2005), though this assumption has been recently jeopardized (Nickstadt *et al*. 2004). Therefore, testing the role of salicylates in our pathosystem would be of interest.

### Root communication also plays a role in the TuMV - *A. thaliana* pathosystem

Addressing our fourth and final question, there is abundant data demonstrating the functions of mycorrhizae and root exudates in supplying between-plant cues and signals (Babikova *et al*. 2013; Wang *et al*. 2020), but there is significantly less recorded evidence of belowground volatile-mediated interplant communication. Our exploratory experiment produced somehow puzzling results. No changes in disease progression have been seen in young receiver plants, regardless of whether they were capable of root-communication or not. Adult receivers, on the other hand, experienced milder symptoms and a more gradual onset of symptoms when allowed for belowground communication. This growth stage-dependent effect could simply result from a slower soil diffusion of signaling molecules, which do not reach receivers when emitters are inoculated concurrently but do so when emitters are inoculated well in advance. Root exudates, like VOCs, contain a complex mixture of compounds, such as ethylene, strigolactones, JA, and allantoin (Wang et al, 2020), as well as microRNAs arranged into extracellular vesicles (Betti et al. 2021). Further research is necessary to determine the chemical nature of this underground signals.

## Conclusions

Our exploratory work suggests a possible role of VOCs, and in particular of JA, in the attenuation of symptoms’ progression in newly infected plants living alongside already infected ones. In this sense, high disease prevalence will result in more and more plants emitting VOCs and in a reduction of susceptibility of noninfected receiver individuals. It is interesting to note that younger plants, which are more sensitive to VOCs signaling than their more mature neighbors, exhibit higher level of protection. Finally, although it merits further investigation, another root communication pathway may also have an impact on this protective community effect.

## Acknowledgements

We thank Paula Agudo for technical assistance and Francesc X. Mulet i Piera, Enric Pèrez Parets and Mario Ruiz Pérez for help. This work was supported by grant PID2019-103998GB-I00 funded by MCIN/AEI/10.13039/501100011033.

## Supplementary Materials

**Table S1.** Results of fitting the severity of symptoms progression data shown in Fig. 1 to a repeated measures ANOVA model.

| Effect | *SS* | *d.f.* | *F* | *P* | $\eta_P^2$ | $1 - \beta$ |
| --- | --- | --- | --- | --- | --- | --- |
| **Within-plant tests<sup>1</sup>** |  |  |  |  |  |  |
| dpi | 861.4954 | 1.9259 | 337.7671 | < 0.0001 | 0.5708 | 1 |
| dpi by prevalence | 8.1168 | 1.9259 | 3.1824 | 0.0442 | 0.0124 | 0.5970 |
| dpi by growth stage | 77.8008 | 1.9259 | 30.5034 | < 0.0001 | 0.1072 | 1 |
| dpi by prevalence by growth stage | 13.9333 | 1.9259 | 5.4628 | 0.0050 | 0.0211 | 0.8376 |
| error within dpi | 647.8425 | 489.1804 |  |  |  |  |
| **Between-plant tests** |  |  |  |  |  |  |
| intersection | 1320.3317 | 1 | 628.5568 | < 0.0001 | 0.7122 | 1 |
| prevalence | 25.5676 | 1 | 12.1717 | 0.0006 | 0.0457 | 0.9352 |
| growth stage | 234.8936 | 1 | 111.8234 | < 0.0001 | 0.3057 | 1 |
| prevalence by growth stage | 0.1211 | 1 | 0.0576 | 0.8104 | 0.0002 | 0.0566 |
| error | 533.5464 |  |  |  |  |  |

<sup>1</sup>Greenhouse-Geisser correction for data sphericity ( $\varepsilon = 0.1751$ ).

**Table S2.** Results of fitting the severity of symptoms progression data shown in Fig. 2 to a repeated measures ANOVA model.

| **Effect** | ***SS*** | ***d.f.*** | ***F*** | ***P*** | **$\eta_p^2$** | **$1 - \beta$** |
| --- | --- | --- | --- | --- | --- | --- |
| **Within-plant tests<sup>1</sup>** |  |  |  |  |  |  |
| dpi | 1311.9225 | 3.3097 | 771.3786 | < 0.0001 | 0.7068 | 1 |
| dpi by prevalence | 8.0036 | 3.3097 | 4.7059 | 0.0020 | 0.0145 | 0.9187 |
| dpi by plant genotype | 7.8633 | 3.3097 | 4.6234 | 0.0023 | 0.0142 | 0.9136 |
| dpi by prevalence by plant genotype | 8.5237 | 3.3097 | 5.0117 | 0.0013 | 0.0154 | 0.9353 |
| error within dpi | 544.2401 |  |  |  |  |  |
| **Between-plant tests** |  |  |  |  |  |  |
| intersection | 2925.9232 | 1 | 2946.1745 | < 0.0001 | 0.9020 | 1 |
| prevalence | 15.0250 | 1 | 15.1290 | 0.0001 | 0.0451 | 0.9724 |
| plant genotype | 17.0618 | 1 | 17.1798 | < 0.0001 | 0.0510 | 0.9851 |
| prevalence by plant genotype | 16.5458 | 1 | 16.6603 | < 0.0001 | 0.0495 | 0.9825 |
| error | 317.8004 | 320 |  |  |  |  |

<sup>1</sup>Greenhouse-Geisser correction for data sphericity ( $\varepsilon = 0.3310$ ).

**Table S3.** Results of fitting the severity of symptoms progression data shown in Fig. 3 to a repeated measures ANOVA model.

| **Effect** | ***SS*** | ***d.f.*** | ***F*** | ***P*** | **$\eta_p^2$** | **$1 - \beta$** |
| --- | --- | --- | --- | --- | --- | --- |
| **Within-plant tests<sup>1</sup>** |  |  |  |  |  |  |
| dpi | 2754.8499 | 3.0090 | 560.8404 | < 0.0001 | 0.7247 | 1 |
| dpi by plants per pot | 17.7831 | 3.0090 | 3.6203 | 0.0129 | 0.0167 | 0.7966 |
| dpi by inoculation delay | 44.5665 | 6.0180 | 4.5365 | 0.0002 | 0.0409 | 0.9872 |
| dpi by plants per pot by inoculation delay | 28.8128 | 3.0090 | 5.8658 | 0.0006 | 0.0268 | 0.9541 |
| error within dpi | 1046.2568 | 640.9190 |  |  |  |  |
| **Between-plant tests** |  |  |  |  |  |  |
| intersection | 7100.0488 | 7 | 1108.2300 | < 0.0001 | 0.8388 | 1 |
| plants per pot | 61.8049 | 1 | 9.6470 | 0.0022 | 0.0433 | 0.8712 |
| inoculation delay | 68.9234 | 2 | 5.3790 | 0.0053 | 0.0481 | 0.8389 |
| plants per pot by inoculation delay | 79.1687 | 1 | 12.3573 | 0.0005 | 0.0548 | 0.9382 |
| error | 1364.6178 | 213 |  |  |  |  |

<sup>1</sup>Greenhouse-Geisser correction for data sphericity ( $\varepsilon = 0.2006$ ).

## Notes

https://doi.org/10.5281/zenodo.7663956

### Competing Interest Statement

The authors have declared no competing interest.

## References

Babikova Z, Gilbert L, Bruce TJ, et al. 2013. Underground signals carried through common mycelial networks warn neighbouring plants of aphid attack. Ecol Lett 16: 835–843.

Baldwin IT, Schultz JC. 1983. Rapid changes in tree leaf chemistry induced by damage: evidence for communication between plants. Science 221: 277–279.

Betti F, Ladera-Carmona M., Weits DA, et al. 2021. Exogenous miRNA induces post-transcriptional gene silencing in plants. Nat Plants 7: 1379–1388.

Bitas V, Kim HS, Bennett JW, et al. 2013. Sniffing on microbes: diverse roles of microbial volatile organic compounds in plant health. Mol Plant Microbe Interac 26: 835–843.

Boyes DC, Zayed AM, Ascenzi R, et al. 2001. Growth stage-based phenotypic analysis of Arabidopsis: a model for high throughput functional genomics in plants. Plant Cell 13: 1499–1510.

Brosset A, Blande JD. 2022. Volatile-mediated plant-plant interactions: volatile organic compounds as modulators of receiver plant defence, growth, and reproduction. J Exp Bot 73: 511–528.

Butković A, González R, Rivarez MPS, et al. 2021. A genome-wide association study identifies Arabidopsis thaliana genes that contribute to differences in the outcome of infection with two turnip mosaic potyvirus strains that differ in their evolutionary history and degree of host specialization. Virus Evol 7: veab063.

Chen CC, Chao CH, Chen CC, et al. 2003. Identification of turnip mosaic virus isolates causing yellow stripe and spot on calla lily. Plant Dis 87: 901–905.

Cohen J. 1973. Eta-squared and partial eta-squared in fixed factor ANOVA designs. Educational and Phsychol Measur 33: 107–112.

Corrêa RL, Sanz-Carbonell A, Kogej Z, et al. 2020. Viral fitness determines the magnitude of transcriptomic and epigenomic reprograming of defense responses. Mol Biol Evol 13: 1866–1881.

Craufurd PQ, Wheeler TR. 2009. Climate change and the flowering time of annual crops. J Exp Bot 60: 2529–2539.

Dolch R, Tscharntke T. 2000. Defoliation of alders (Alnus glutinosa) affects herbivory by leaf beetles on undamaged neighbours. Oecologia 125: 504–511.

Falik O, Mauda S, Novoplansky A. 2023. The ecological implications of interplant drought cuing. J Ecol 111: 23–32.

Gao S, Dai X, Wang L, et al. 2022. Epidemic spreading in metapopulation networks coupled with awareness propagation. IEEE Trans Cybernetics.

García-Ruiz H, Murphy JF. 2001. Age-related resistance in bell pepper to cucumber mosaic virus. Ann Appl Biol 139: 307–317.

Ghosh S, Didi-Cohen S, Cna’ani A, et al. 2022. Comparative analysis of volatiles emitted from tomato and pepper plants in response to infection by two whitefly-transmitted persistent viruses. Insects 13: 840.

Girden ER. 1992. ANOVA: Repeated Measures. Sage Publications Inc, London, UK, 84 pp.

Glazebrook J. 2005. Contrasting mechanisms of defense against biotrophic and necrotrophic pathogens. Annu Rev Phytopathol 43: 205–227.

González R, Butković A, Elena SF. 2019. Role of host genetic diversity for susceptibility-to-infection in the evolution of virulence of a plant virus. Virus Evol 5: vez024.

Granell C, Gómez S, Arenas A. 2013. Dynamical interplay between awareness and epidemic spreading in multiplex networks. Phys Rev Lett 111: 128701.

Greenhouse SW, Geisser, S. 1959. On methods in the analysis of profile data. Psychometrika 24: 95–112.

Hammerbacher A, Coutinho TA, Gershenzon J. 2019. Roles of plant volatiles in defence against microbial pathogens and microbial exploitation of volatiles. Plant Cell Environ 42: 2827–2843.

Hu L, Yang L. 2019. Time to fight: molecular mechanisms of age-related resistance. Phytopathology 109: 1500–1508.

Huang Y, Hong H, Xu M, et al. 2020. Developmentally regulated Arabidopsis thaliana susceptibility to tomato spotted wilt virus infection. Mol Plant Pathol 21: 985–998.

Islam W, Naveed H, Zaynab M, et al. 2019. Plant defense against virus diseases; growth hormones in highlights. Plant Signal Behav 14: e15967189.

Kus JV, Zaton K, Sarkar R, et al. 2002. Age-related resistance in Arabidopsis is a developmentally regulated defense response to Pseudomonas syringae. Plant Cell 14: 479–490.

Laurie-Berry N, Joardar V, Street IH, et al. 2006. The Arabidopsis thaliana JASMONATE INSENSITIVE 1 gene is required for suppression of salicylic acid-dependent defenses during infection by Pseudomonas syringae. Mol Plant Microbe Interac 19: 789–800.

Lee K, Seo PJ. 2014. Airborne signals from salt-stressed Arabidopsis plants trigger salinity tolerance in neighboring plants. Plant Signal Behav 9: e28392.

Levy D, Lapidot M. 2008. Effect of plant age at inoculation on expression of gene resistance to tomato yellow leaf curl virus. Arch Virol 153: 171–179.

Li T, Xiao Y. 2021. Linking the disease transmission to information dissemination dynamics: an insight from a multi-scale model study. J Theor Biol 526: 110796.

Loreto F, D’Auria S. 2022. How do plants sense volatiles sent by other plants? Trends Plant Sci 27: 29–38.

Melero I, González R, Elena SF. 2023. Host developmental stages shape the evolution of a plant RNA virus. Phil Trans R Soc B 378: 20220005.

Moreira X, Granjel RR, de la Fuente M, et al. 2020. Plant Cell Environ 44: 1192–1201.

Navarro R, Ambrós S, Butković A, et al. 2022. Defects in plant immunity modulate the rates and patterns of RNA virus evolution. Virus Evol 8: veac059.

Neil K, Wu J. 2006. Effects of urbanization on plant flowering phenology: a review. Urban Ecosyst 9: 243–257.

Nickstadt A, Thomma BPHJ, Feussner I, et al. 2004. The Jasmonate-insensitive mutant jin1 shows increased resistance to biotrophic as well as necrotrophic pathogens. Mol Plant Pathol 5: 425–434.

Peñuelas J, Llusià J. 2004. Plant VOC emissions: making use of the unavoidable. Trends Ecol Evol 19: 402–404.

Riedlmeier M, Ghirardo A, Wening M, et al. 2017. Monoterpens support systemic acquired resistance within and between plants. Plant Cell 29: 1440–1459.

Santonja M, Bousquet-Mélou A, Greff S, et al. 2019. Allelopathic effects of volatile organic compounds released from Pinus halepensis needles and roots. Ecol Evol 9: 8201–8213.

Scatà M, Di Stefano A, Liò P, et al. 2016. The impact of heterogeneity and awareness in modeling epidemic spreading on multiplex networks. Sci Rep 6: 37105.

Shulaev V, Silverman P, Raskin I. 1997. Airborne signalling by methyl salicylate in plant pathogen resistance. Nature 385: 718–721.

Tamogami S, Rakwal R, Agrawal G.K. 2008. Interplant communication: airborne methyl jasmonate is essentially converted into JA and JA-Ile activating jasmonate signaling pathway and VOCs emission. Biochem Biophys Res Commun 376: 723–727.

Tun W, Yoon J, Jeon JS, et al. 2021. Influence of climate change on flowering time. J Plant Biol 64: 193–203.

Ueda H, Kikuta Y, Matsuda K. 2012. Plant communication: mediated by individual or blended VOCs? Plant Signal Behav 7: 222–226.

Wang NQ, Kong CH, Wang P, et al. 2020. Root exudate signals in plant-plant interactions. Plant Cell Environ 44: 1044–1058.

Wang Z, Andrews MA, Wu ZX, et al. 2015. Coupled disease-behavior dynamics on complex networks: a review. Phys Life Rev 15: 1–29.

Wilcoxon F. 1945. Individual comparisons by ranking methods. Biomet Bull 1: 80–83.

Wolf AA, Zavaleta ES, Selmants PC. 2017. Flowering phenology shits in response to biodiversity loss. Proc Natl Acad Sci USA 114: 3463–3468.

